# Climate and land use change variables affect microbial assemblage and denitrification capability in organic-rich subterranean estuaries

**DOI:** 10.1101/2023.06.23.546288

**Authors:** Dini Adyasari, Natasha T. Dimova, Sinead Ni Chadhain, Hannelore Waska

## Abstract

Microbial communities in subterranean estuaries (STE) mediate biogeochemical reactions of coastal groundwater discharging to the oceans; however, studies on their response to abrupt environmental changes caused by climate and land use changes are still limited. In this study, we conducted a controlled laboratory study using combined geochemical and metagenomic approaches to investigate microbial structures and their metabolic pathways under different nitrate (NO_3_^−^) inputs, saline solution, and incubation times, which were used as proxies of land use, salinization of the shallow aquifer, and climate changes. We found a highly reducing habitat and amplification of genes related to denitrification, sulfate reduction, and methanogenesis processes. Core communities consisted of Clostridia, Bacilli, Alphaproteobacteria, Gammaproteobacteria, and Desulfobaccia were observed. The qualitative degradation of terrestrial, plant-derived organic matter (i.e., tannin and lignin) was predicted to not being affected by environmental changes because of it being implemented by core communities and the abundance of electron donor and acceptors. We observed that the assemblages of less prevalent taxa were influenced by seasonal sampling and incubation times, while denitrification was affected by groundwater and seawater inputs. Long-term incubation gave sufficient time for microbes to degrade less labile DOM, promoted the re-release of buried solid phase organic matter into the active carbon cycle, and increased the relative abundance of biofilm or spore-forming taxa while decreasing that of rare taxa such as methanogenic archaea. Our results illustrate the sensitivity of microbial assemblages to environmental change and their capacity to mediate C and N cycles in coastal areas, further affecting coastal water quality and ecosystem-scale biogeochemistry.

**Plain Language Summary:** This study investigated how microbial communities in subterranean estuaries (STE) respond to climate and land use changes. Understanding microbial responses is essential, considering they control the degradation of terrestrial solutes transported to the ocean. STE sediments collected from different seasons were incubated with different nitrate inputs, saline solution, and incubation times to represent changing groundwater quality, sea level rise, and groundwater residence time, respectively. The relative proportions of core microbial groups (Clostridia, Bacilli, Alphaproteobacteria, Gammaproteobacteria, and Desulfobaccia) were stable across all treatments; however, less adaptable groups did not survive long incubation times. Seawater addition negatively affected nitrate removal, while plant-derived organic matter degradation was not significantly influenced by changing environmental parameters. The study highlights how microbial communities and metabolic processes related tothe carbon and nitrogen cycles are susceptible to environmental change. Ultimately, these changes in the microbial community can affect water quality and ecosystem health in coastal areas. This study investigated how microbial communities in subterranean estuaries (STE) relative proportions of core microbial groups (Clostridia, Bacilli, Alphaproteobacteria, Gammaproteobacteria, and Desulfobaccia) were stable across all treatments; health in coastal areas.

**Key points:**

- Core communities’ proportions were stable across different treatments and contributed to plant-derived DOM degradation alongside fermenters and methanogens.
- Sediment denitrification capability was associated with groundwater and seawater input.
- Long-term groundwater residence time negatively influenced rare biosphere.

## 1. Introduction

Subterranean estuaries (STE), the mixing zone between meteoric groundwater and seawater in shallow coastal aquifers, are known to be hotspots for biogeochemical reactions in coastal areas (Moore, 1999). The biogeochemical processes and the export of STE products influence the quality of receiving coastal water and ecosystems (Moosdorf et al., 2021; Rocha et al., 2021). For instance, groundwater discharge discharging from STEs has been recognized to contribute to algal bloom proliferation, bottom water hypoxia, and ocean acidification in marine water (Dulai et al., 2021; Montiel et al., 2019b; Santos et al., 2021). Several factors are known to modulate the biogeochemical STE processes, such as redox conditions (Slomp and Van Cappellen, 2004), residence time (Gonneea and Charette, 2014), groundwater-seawater mixing rate (Kroeger and Charette, 2008), and the availability of electron donors and acceptors (Couturier et al., 2017).

In STEs, oxidants are consumed in order of decreasing energy production per mole of organic carbon (OC): oxygen > nitrate (NO_3_^−^) > manganese oxides (MnO_2_) > ferric iron (Fe^3+^) > sulfate (SO_4_^2−^) (Froelich et al., 1979). In coastal areas with high anthropogenic activities, terrestrial groundwater often delivers NO_3_^−^ to STE, where it is used as the primary electron acceptor in denitrification process or preserved as nitrogen (N) in the system by dissimilatory nitrate reduction to ammonium (DNRA) (Adyasari et al., 2020; Calvo-Martin et al., 2022; Jiao et al., 2018; Sáenz et al., 2012; Santoro, 2010). In deep, more anaerobic STEs, biogeochemical transformations are commonly mediated by sulfate-reducing bacteria (SRB) and methanogenic archaea, and consequently, accumulate sulfide and methane (CH_4_) in these layers (Hong et al., 2019; Purkamo et al., 2022).

Most of microbial respiration in STEs require organic or carbon (C) substrates, which can be found autochthonously from terrestrial plant material or allochthonous from marine sources (Seidel et al., 2015). Terrestrial-based dissolved organic matter (DOM) is often dominated by aromatic and highly unsaturated compounds and originated from buried or decomposed plant bark, leaves, and seeds (Koch and Dittmar, 2006; Stenson et al., 2003). Such compounds include tannin and lignin, the main component of plant biomass. Considering their highly refractory characteristics, these compounds have previously not been considered an important energy source for subsurface communities. Yet, the capability to degrade these resistant organic compounds into metabolites and final products (e.g., gallate, methyl gallate, catechol, protocatechuic acid, vanillin, vanillic acid) have been observed in different fungi and bacterial strains (Bhat et al., 1998; Bugg et al., 2011). For instance, lignin-degrading genes are widespread among taxonomic groups of Actinobacteria, Gammaproteobacteria, Alphaproteobacteria, and Cyanobacteria (Bugg et al., 2011). In recent years, studies combining 16S rRNA sequencing and DOM molecular characterization by Fourier Transform Ion Cyclotron Resonance Mass Spectrometry (FTICRMS) have linked decarboxylation of tannin- and lignin-like structures to heterotrophic bacteria and candidate phyla radiation group in coastal sediments (Degenhardt et al., 2021; Oni et al., 2015).

Because of their role in ecosystem functions, understanding microbial roles and their reaction to STE dynamics is particularly interesting. Even though studies focusing on the microbial aspect of STE have gained attention in recent years, there is still lack of data in this field compared to research related to the physicochemical characteristics of STE (Ruiz-González et al., 2021). This can be attributed to the highly heterogenous nature of STEs, which further leads to difficulty linking microbial groups to metabolic activities. For instance, microbial communities are known to be highly diverse both in (1) spatial distribution, as a result of steep physicochemical gradient (Adyasari et al., 2019; Ruiz-González et al., 2022), and (2) temporal distribution, due to rapid and fluctuating changes of their surrounding habitats, such as tidal cycles (Moosdorf et al., 2021; Retter et al., 2021). In so-called “hot moments” of STE, temporal equivalents of “hotspot”, coastal groundwater chemical and biological compositions may be altered on short timescales, e.g., in the span of hours during different tidal cycles (Lee et al., 2017). With that being said, other STE studies also observed the occurrence of core microbial communities in STEs, i.e., a resilient group of microorganisms usually comprises of generalists or cosmopolitan taxa that exhibit lack of temporal or spatial variability (Calvo-Martin et al., 2022; Degenhardt et al., 2020). However, due to scarcity of such studies, a particular attention is still needed as environmental alteration caused by climate (sea level rise; intensified precipitation, land use (increasing pollutant [e.g., NO_3_^−^, PO_4_^3-^] concentration in the aquifer), and water usage (groundwater withdrawal) changes, are forecasted to create disturbance to coastal microbial communities and ecosystem functions they serve (Retter et al., 2021).

To elucidate the impact of extreme hydrological and meteorological changes on the structure and metabolic activity of the microbial community, we collected STE sediments from a site in the northeastern Gulf of Mexico and subjected them to controlled laboratory experiments. The Gulf of Mexico is historically characterized by hypoxic events (Turner et al., 1987), a high frequency of hurricanes (Trepanier et al., 2015), and coastal aquifer salinization in recent decades (Lin et al., 2009). We targeted STE depths up to 2-3 m, which were previously identified as the depth of shallow groundwater flow in the study site (Montiel et al., 2019b). The microcosm study was implemented using sediments recovered during different seasons and incubated with a wide range of NO_3_^−^ concentrations, saline solution, and incubation times—which were used as proxies of climate, land use, salinization of the shallow aquifer, and water usage changes mentioned above. Using a multifaceted method approach of geochemical measurement, FTICRMS, quantitative Polymerase Chain Reaction (qPCR), and 16S rRNA gene sequencing, we aim to investigate identify taxonomic groups mediating terrestrial organic matter degradation and their metabolic role in C and N cycles.

## 2. Materials and Methods

### 2.1 Study site

The shallow aquifer in Mobile Bay, located in the northeastern Gulf of Mexico, is up to 3 m deep and comprised of a up to 1.5-m organic-rich, black fine sand discontinuous layer (Montiel et al., 2019a). The organic layer was formed around the 1960s and is rich in Fe (Lamore, 2020). DOM characterization using optical markers suggested that this layer consists primarily of humic-like components C2 and C1, originating from decomposing salt marsh plants such as *Juncus roemerianus, Spartina alterniflora*, and freshwater marsh *Typha* and *Schoenoplectus* (Montiel et al., 2019b). The STE through which terrestrial groundwater percolates is mostly hypoxic; as a result, submarine groundwater discharge (SGD) transports high ammonium (NH_4_^+^) and dissolved organic nitrogen (DON) concentrations to coastal water, further creating bottom water hypoxia (Montiel et al., 2019a; Montiel et al., 2019b). A previous field study at this site revealed highly diverse vertical stratification of porewater microbial communities, ranging from Cyanobacteria in the upper layer and anaerobic bacteria, SRB, fermenter, and methanogenic archaea in the middle and bottom layers (Adyasari et al., 2020).

### 2.2 Experimental incubations and physicochemical analysis

Sediment cores were recovered during contrasting seasons. The first sediment collection was conducted during the storm season in April 2021, and the second one during normal-dry conditions (September 2021). The cores were recovered from a 2 m^2^ plot of the intertidal zones using a GeoProbe model 5410 direct push technology (DPT). Upon collection, the sediment cores were transported within 2-3 days to the Coastal Hydrogeology Laboratory at the University of Alabama and refrigerated before analyses and experiments. The sediment cores were sliced into 10 cm sections and evaluated for their organic matter content following ASTM methods 2974-87 (American Society of Testing Materials, 1993). Sections with more than 10% organic content were chosen for the incubation experiment and thoroughly mixed and homogenized. From each season, duplicate aliquots of 250 mg of sediments were set apart for particle size measurement and sedimentary microbial characterization before incubation (i.e., samples O-Storm-1, O-Storm-2, O-Dry). The particle size distribution analysis was conducted using laser particle sizer analyzer. Briefly, 10 grams of homogenized sediments were dried and pre-treated with sodium hexametaphosphate before undergoing laser diffraction and imaging in a Bettersizer S3 Plus (Bettersizer Instruments Ltd., China).

We conducted sediment incubations with four setups as visualized in Figure 1 and described as the following:

1. *Seasonal dynamics (storm vs. dry season)*. To examine the seasonal impacts on microbial community composition, we collected STE sediments in two contrasting sampling sessions. Since the sampling campaign in April was conducted during the storm season, they were labeled as “Storm” batch samples. Meanwhile, incubations conducted with sediments collected in September were named “Dry” batch samples.
2. *NO_3_^−^ inputs.* To elucidate microbial response to changing groundwater quality, sediment samples were incubated with a wide range of NO_3_^−^ concentrations: 50 μM (Storm-2 and Dry-2), 100 μM (Storm-3 and Dry-3), and 200 μM of an artificial NO_3_^−^ solution (Storm-4 and Dry-4), and groundwater collected from the field (NO_3_^−^ concentration of 210 μM, Dry-GW-1 and Dry-GW-2). Groundwater was collected from a 2 m-deep coastal well and stored in a refrigerator before it was used for the experiment. Note that groundwater addition to the incubation was only conducted during Dry batch samples, as we could not obtain groundwater samples during the storm season. The artificial NO_3_^−^ solution was prepared using Nanopure water and high-purity grade sodium nitrate (VWR Chemicals).
3. *Saline solution.* To investigate the impact of STE salinization on microbial communities, sediment samples were incubated with different saline solution: NaCl solution with 50 μM NO_3_^−^ and salinity = 10 (Storm-5 and Dry-5), NaCl solution with a 50 μM NO_3_^−^ and salinity = 20 (Storm-6 and Dry-6), and coastal seawater collected from Mobile Bay with 5 μM NO_3_^−^ and salinity = 15 (Dry-SW-1 and Dry-SW-2). Seawater was collected from 0.2 m depth of Mobile Bay water column and stored in refrigerator prior to its usage in the experiment. Like the freshwater setup, incubations with seawater from the field were only supplemented with sediments from the dry season. The artificial NaCl solution was prepared by mixing Nanopure water with combusted pure NaCl salt (VWR chemicals).
4. *Incubation times.* As a final variable, we incubated sediments over different timescales, mimicking scenarios of altered STE residence time. In this experimental setup, all batch samples were incubated for different periods, i.e., 5 days (Storm-2 and Dry-2), 10 days (Storm-7 and Dry-7), 15 days (Storm-8 and Dry-8), 20 days (Storm-9 and Dry-9), 25 days (Storm-10), 30 days (Storm-11), and 35 days (Storm-12). The 5-day incubation was employed to simulate the groundwater residence time previously reported for the local STE (Montiel et al., 2019a). We used 50 μM of NO_3_^−^ solution with zero salinity as input for this set of experiments.

**Figure 1.**
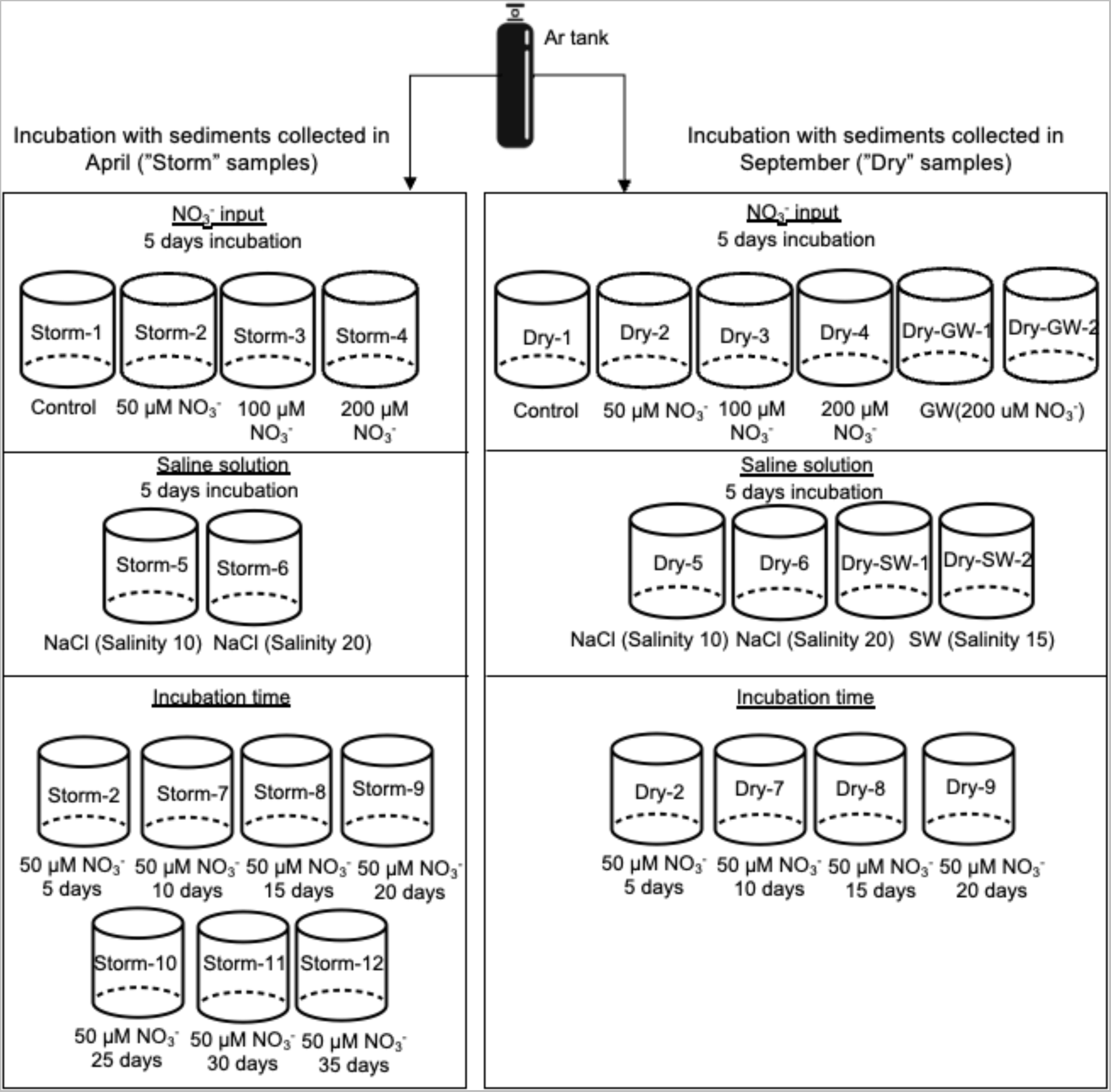
Schematic diagram of the incubation experiments. GW and SW refer to groundwater and seawater collected from the site, respectively.

In all treatments, Nanopure water with zero salinity and zero NO_3_^−^ concentration were used for the control samples (Storm-1 and Dry-1). The glass jars were sterilized by autoclave (121°C for 20 min) prior to their usage for the experiment. The slurries for the experiments were made by mixing of solution and sediments with a 1:1 ratio and brought to hypoxic conditions (dissolved oxygen (DO) < 2 mg L^−1^)) by purging them with ultra-pure-grade argon gas. The incubations were conducted inside an oxygen-depleted chamber (ODC). DO levels within the ODC were monitored continuously using an atmospheric oxygen sensor (Vernier, Accuracy ± 1%); oxygen levels were always < 2 mg L^−1^. The jars were kept in the dark to mimic the subsurface conditions. After incubation, porewater samples were collected to determine physical parameters (e.g., temperature, DO, salinity, pH), NO_3_^−^, NH_4_^+^, phosphate (PO_4_^3^)^−^, dissolved organic carbon (DOC), DON, and DOM. Physical parameters were measured directly in the laboratory using a pre-calibrated YSI Pro 2030 handheld instrument. Aliquots for nutrient analyses were filtered through a 0.45 μm cellulose acetate filter, frozen, and sent to the analytical facilities at Dauphin Island Sea Laboratory (DISL). Aliquots for DOC, DOM, and total dissolved nitrogen (TDN) analyses were filtered through a 0.22 μm polyethersulfone filter, acidified with HCl to pH = 2, and kept at 4°C temperature until further analysis in the ICBM-MPI Bridging Group Marine Geochemistry laboratory at the University of Oldenburg, Germany. DON concentration was calculated by subtracting TDN and dissolved inorganic nitrogen (e.g., NO_3_^−^, NO_2_^−^, NH_4_^+^). The results from the incubations are presented in Table S1.

DOM samples were extracted and analyzed with FTICRMS according to Waska et al. (2021). The ICBM-OCEAN tool was used to assess chemical characteristics (elemental ratios and compound classes [highly unsaturated [Hu], aromatic [Ar], unsaturated [Us], and saturated [Sat]) and molecular indices (i.e., aromaticity index [AI.mod] (Koch and Dittmar, 2006), lability index [MLB_l] (D’Andrilli et al., 2015), degradation index [I_deg_] (Flerus et al., 2012), terrestrial index [I_Terr_] (Medeiros et al., 2016), and biological productivity index [I_bioprod_]) (Rossel et al., 2020). According to Merder et al. (2020), highly unsaturated and aromatic compounds are categorized as “less labile” compounds, while “labile” compounds consist of unsaturated and saturated compounds. The complete list of DOM compounds identified by FTICRMS can be found in Table S2.

### 2.3 DNA extraction, amplification, and sequencing

250 mg of original and treated sediments from each incubation jar were collected for sequencing analysis. DNA extraction was performed using the DNeasy® PowerSoil (Qiagen, USA) as specified by the manufacturer. DNA pellets were sent to Michigan State University Genomic Laboratory for paired-end amplicon sequencing on an Illumina MiSeq platform. DNA sequences of the V3-V4 hypervariable region of the 16s rRNA gene were obtained with primer set pair 515F/806R (Caporaso et al., 2011). The sequences provided by Michigan State University were processed using DADA2 version 1.24.0 (Callahan et al., 2016) on R version 4.2.1. Taxonomic assignment of the different amplicon sequence variance (ASV) was performed against SILVA version 138.1. Unwanted lineages (chloroplasts, mitochondria, and ASVs unclassified on phylum level) were removed from the dataset. Because of insufficient sequencing depth, samples Dry-5 and O-Dry-2 were not used for further downstream analyses. The complete ASV and taxonomic table can be found in Table S3. All raw sequences used in this study are publicly available at the NCBI Sequence Read Archive under project accession number PRJNA985793.

### 2.4 PCR and qPCR assays

PCR assays of anaerobic N cycles (denitrification, DNRA, anaerobic ammonium oxidation [anammox]), sulfate reduction, and methanogenesis were performed to amplify genes mediating such processes in sediment communities. Aliquots of a 20 μL PCR reaction included 10 μL Go-Taq master mix (Promega, USA), 2 μL of each forward and reverse primer, 1 μL template DNA and 5 μL nuclease-free H_2_O. The following biogeochemical processes and reference primers were assessed by PCR assay: denitrification (Throbäck et al., 2004), DNRA (Mohan et al., 2004), anammox (Lam et al., 2009), sulfate reduction (Kondo et al., 2008), and methanogenesis (Luton et al., 2002). The PCR products were purified using the Wizard Sv Gel and PCR Extraction System (Promega) and quantified by Qubit fluorometer (Thermo Fisher Scientific, USA). PCR products were cloned into pCR4-TOPO (Invitrogen, USA) and sequenced prior to use as standards in qPCR.

Further qPCR assay was implemented to investigate denitrification and DNRA gene expression. The reaction setups were prepared by mixing 10 μL of master mix (Thermo Scientific DyNAmo HS SYBR Green qPCR Kits, 2 μL of each forward and reverse primer, 1 μL template DNA, 0.4 μL ROX passive reference dye, and 4.6 μL nuclease-free H_2_O. The qPCR measurement was performed using the Mx3000P qPCR system (Agilent Technologies). Further details concerning PCR and qPCR conditions can be found in Table S4.

### 2.5 Statistical analyses

Microbial results were analyzed in R version 4.2.1 using the vegan package version 2.6.2 (Oksanen et al., 2013). The ASV table was randomly subsampled down to the minimum number of reads per sample to quantify alpha diversity indices (number of ASV [nASV], Shannon diversity index, and the rare biosphere as single sequence OTU [SSO]). Non-metric multidimensional scaling (NMDS) was employed to visualize community composition based on Bray-Curtis dissimilarity, while observed environmental parameters were fitted onto the NMDS using *envfit*. Spearman correlations were used to explore the linkage between microbial classes with DOM molecular compounds from FTICRMS results. The DOM molecular composition table was standardized using centered-log ratio transformation prior to performing Spearman analysis. The p-values were corrected for multiple testing based on the Benjamini-Hochberg method (Benjamini and Hochberg, 1995). Only significant correlations (p-value < 0.05 and Spearman ρ > 0.5 and ρ < −0.5) are presented.

### 2.6 Metabolic prediction

The prediction of functional genes based on 16S rRNA gene sequences was implemented using PICRUST2 (Douglas et al., 2020) via Nephele (Weber et al., 2018). As Mobile Bay STE porewater contained a high concentration of DON, NH_4_^+^, and DOM pool consisting of terrigenous, plant material compounds (Montiel et al., 2019b), we focused on catabolic pathways related to plant-derived aromatic, highly unsaturated compounds, and DON degradation processes. Such processes included catabolic products of tannin and lignin degradation, i.e., gallate, methyl gallate, catechol, vanillin, vanillate, and protocatechuate (Bhat et al., 1998; Sharma et al., 1998). Considering the highly reducing condition in Mobile Bay STE, we also screened metabolic pathways related to sulfur, methane, and fermentation activities. The entire list of pathways generated from the PICRUST2 pipeline can be found in Table S5.

## 3. Results

### 3.1 Physicochemical changes after incubations

In this study, particle size analysis was implemented for homogenized sediments in each season. The results suggested that the average Storm sediment samples contained 6% clay, 44% silt, and 50% sand. In contrast, sediments collected during the dry season had a higher proportion of fine sediments, i.e., 10% clay, 53% silt, and 37% sand. Even though there was a slight variability of particle size distribution, a further multivariate analysis of variance indicated no significant difference between sediments collected in April and September (p > 0.05).

A total of 27 porewater samples were obtained from incubation with diverse sediment collection times, NO_3_^−^ inputs, saline solutions, and incubation times. All samples were depleted of oxygen (average DO = < 3.5 mg L^−1^), had low to moderate pH (ranging from 6.3 to 7.5), and were enriched in DOC and DON, as well as their mineralization products, such as NH_4_^+^ and PO_4_^3-^ concentrations. NO_3_^−^ was removed from porewater with 88-98% removal compared to its concentration prior to incubation. DOM molecular characteristics indicated a higher relative proportion of less labile DOM pools (highly unsaturated and aromatic compounds, average 42%) than labile DOM compounds (unsaturated and saturated compounds, average *7*%).

### 3.2 Microbial diversity and community composition

In total, we observed 65,355 ASVs from 30 samples (three original unmanipulated sediments and 27 incubated sediments). Sediments collected during the storm season exhibited more ASVs and diversity but a similar quantity of rare taxa than samples from the dry season (Figure 2). Artificial NO_3_^−^ solution and groundwater used as inputs to incubation affected alpha diversity in these samples differently: groundwater positively influenced rare taxa but negatively affected the richness of the samples. Artificial NaCl and seawater treatment positively impacted nASV but had less impact on diversity and the number of rare taxa. The long-term incubations had a positive association with nASV and diversity, while significantly reducing the number of rare taxa.

**Figure 2.**
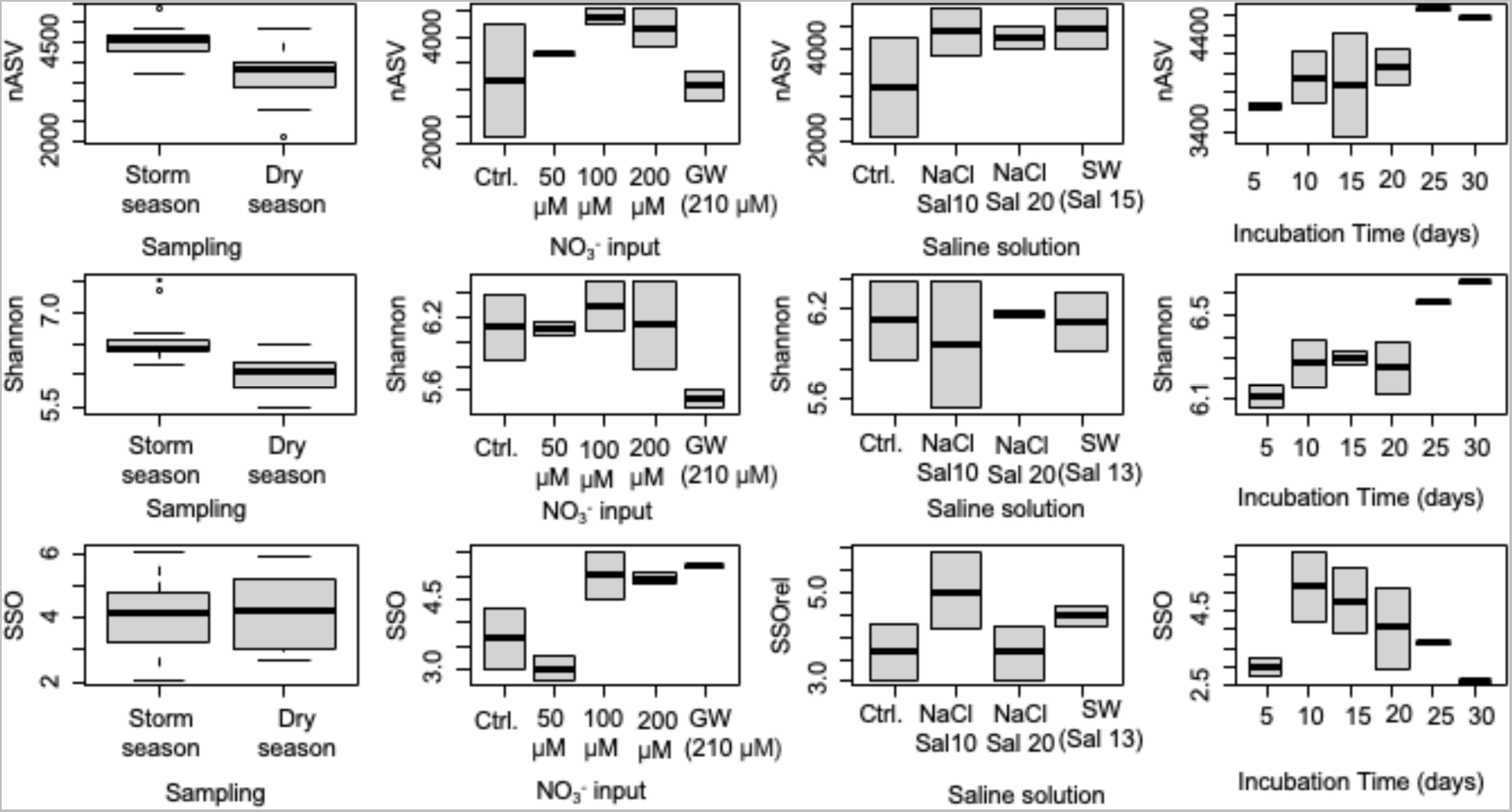
Alpha diversity of samples across different collection times, NO_3_^−^ inputs, saline solutions, and incubation times.

The NMDS plot showed that the highest Bray-Curtis dissimilarity between all samples was observed between incubated sediments collected in different seasons (Figure 3, Bray-Curtis = 0.5). Microbial communities from sediments collected during the storm season were associated with DON concentration and labile, freshly produced organic matter as indicated by MLB_l, unsaturated DOM, and I_bioprod_. In contrast, communities from the dry season were positively correlated with DOC concentrations. Microbial communities in samples with higher salinity were associated with highly unsaturated DOM and NH_4_^+^ and more similar to each other than those found in sediment incubated with zero salinity solution.

**Figure 3.**
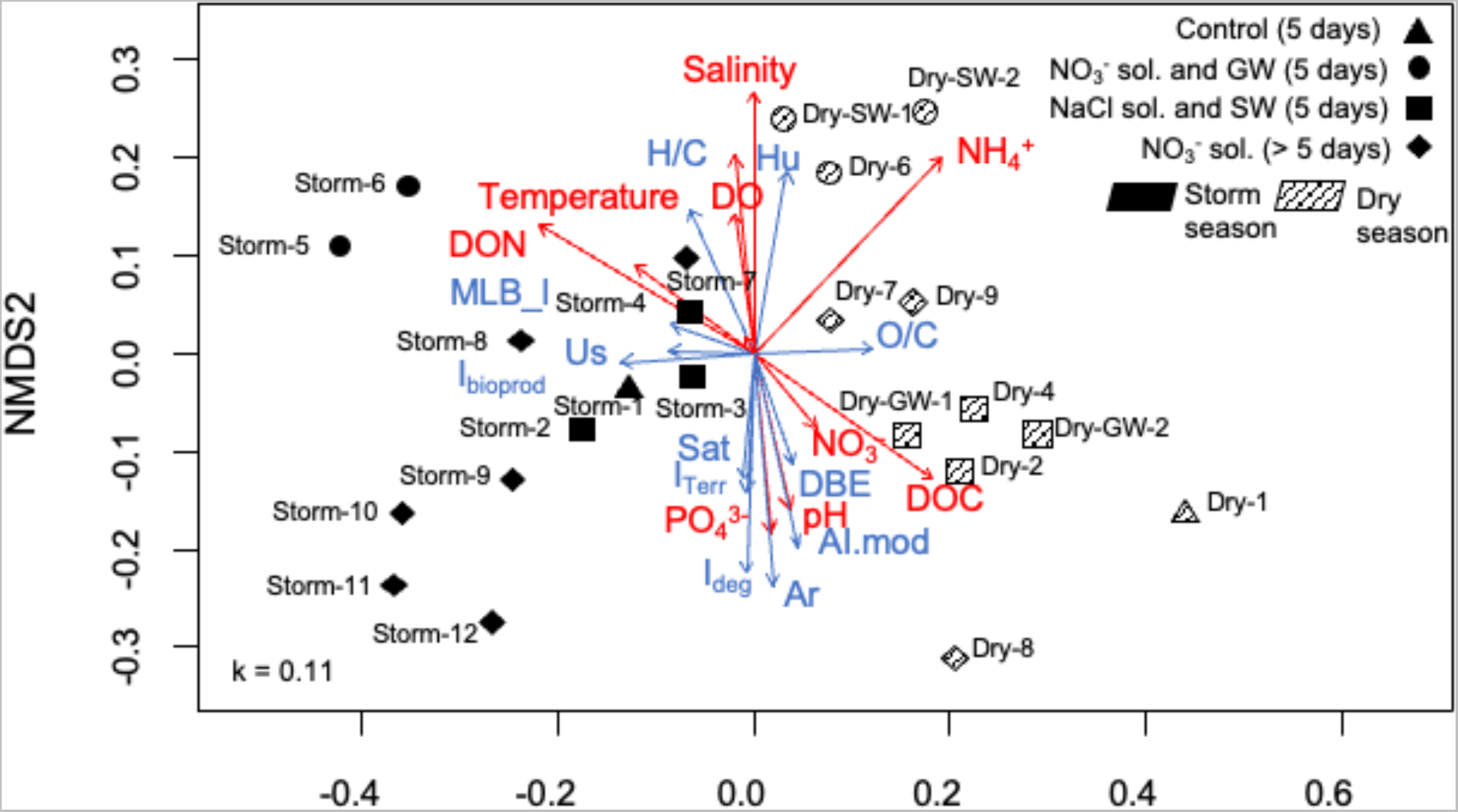
Non-metric multidimensional scaling (NMDS) plot depicting the association between microbial community composition and environmental data across all samples. Red arrows indicate the association of physicochemical parameters with microbial samples, while blue arrows show specific DOM molecular characteristics (indices, elemental ratios, aromaticity) with different microbial samples. SW = seawater, GW = groundwater, Ar = aromatic DOM, Hu = highly unsaturated DOM, Sat = saturated DOM, Us = unsaturated DOM.

The identified bacteria and archaea were all microaerophilic or anaerobic, which coincided with the reduced conditions in the jars and ODC chamber. At the kingdom level, bacteria dominated over archaea across all samples, with archaea comprising between 0.1-1% of the microbial communities. Methanogenic archaea (e.g., Methanosarcinia, Methanocella, Methanobacteria) dominated the samples, while uncultivated Bathyarchaea were also observed (Table S3). In terms of bacteria, we identified a total of 34 phyla across all samples. The core communities included ASVs from Clostridia, Bacilli, Alphaproteobacteria, Gammaproteobacteria, and Desulfobaccia. Clostridia and Bacilli, both belonging to phylum Firmicutes, appeared in all samples with an average of 39% and 29%, respectively (Figure 4). Dominant genera from these two classes detected in all samples were *Paraclostridium, Clostridium sensu stricto*, and *Tepidibacillus* (Figure 4). Alphaproteobacteria (6%, with *Methyloceanobacter* as the most prominent genus) was also identified in all samples, followed by Gammaproteobacteria (average 5%, with genera *Acinetobacter* and *Methylobacter* as the most prominent genera) and Desulfobaccia (4%, with *Desulfobacca* as the most prominent genus). Iron reducers, a group often detected in reducing habitat, were observed with a lower relative proportion than SRB, i.e., between 1-2%. This group was primarily represented by ASVs belonging to Geobacteraceae.

**Figure 4.**
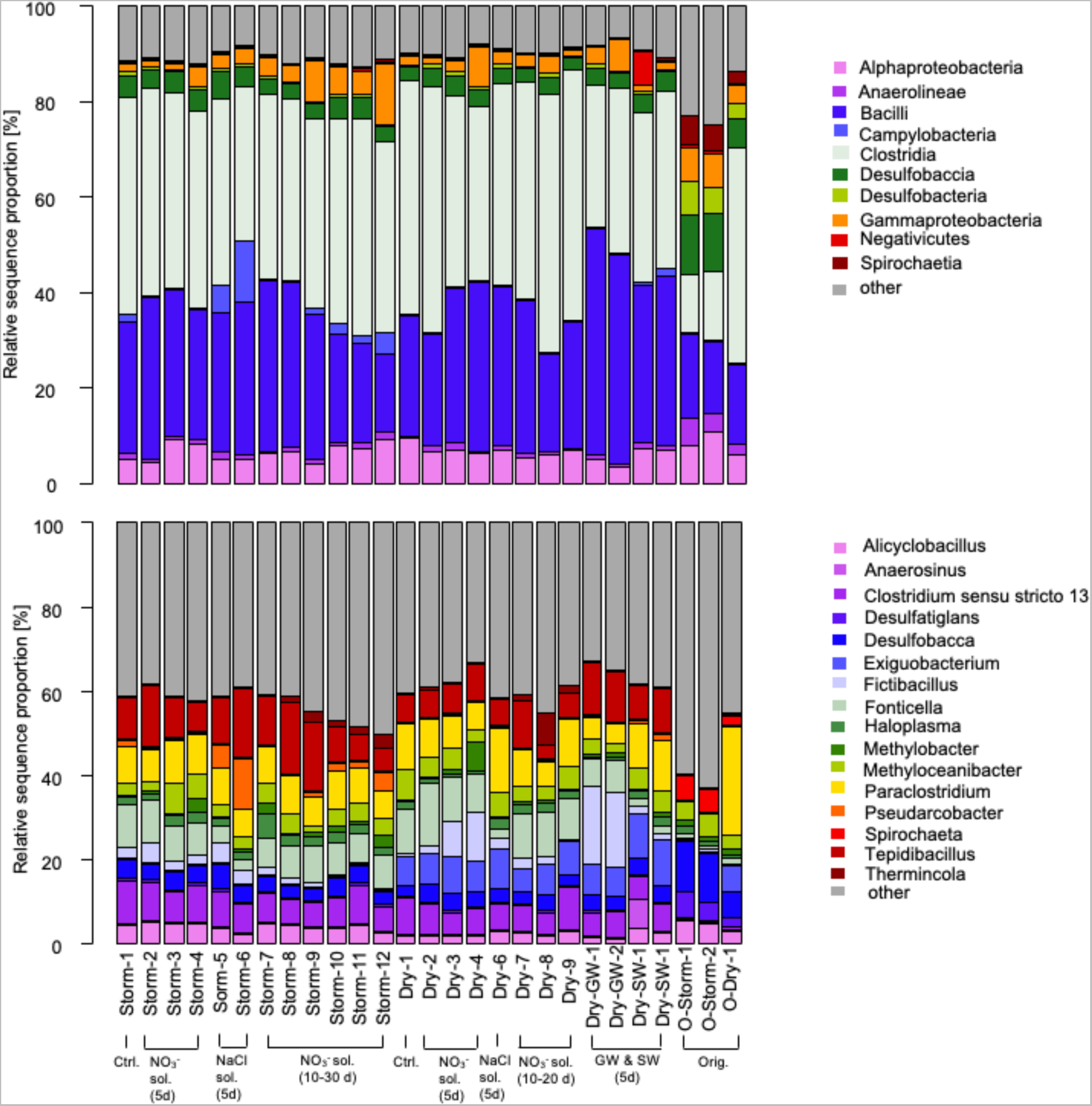
Relative sequence proportion of dominant microbial taxa at class (top) and genus level (bottom).

The original unmanipulated sediments had a similar taxonomic profile as treated sediments with a few notable differences. The relative proportion of Clostridia and Bacilli doubled after incubation (from 16-23% to 30-40%), while Spirochaeta, Desulfobacteria, and Desulfobaccia all decreased in abundance after incubation. Leptospirae (g. *RBG-16-49-21*) and Omnitrophia (g. *Candidatus Omnitrophus*) were microbial classes detected in the original sediments but not in any of the treated samples.

While the core communities remained in stable proportions across all samples, other less prevalent taxa were found to be more prone to respond to physicochemical changes. Campylobacteria (g. *Pseudarcobacter*) were prevalent in sediments collected during storm season (Figure 4). They had a relative proportion in the range of 4% in Storm samples and less than 1% in Dry samples. In contrast, ASVs belonging to the Clostridial genus *Exiguobacterium* were identified in moderate proportions in Dry samples but significantly lower in Storm samples (Figure 4).

Incubation under different NO_3_-concentration settings did not produce changes in microbial community composition. At the same time, sediments from groundwater-spiked samples consisted of a considerable proportion of *Fictibacillus* compared to other samples. Samples with higher salinities (i.e., NaCl solution and seawater) generally displayed distinct microbial community compositions that differentiated them in the NMDS plot (Figure 3). Furthermore, analysis from seawater-spiked incubations showed that Acidimicrobiia (g. *CL500-29 marine group* and *Ilumatobacter)*, Negativicutes (g. *Anaerosinus*), and Cyanobacteria were detected exclusively in these samples.

Microbial communities extracted from sediments treated with longer incubation times (> 5 days) exhibited similar phylogeny to the communities in 5 days incubation; however, some taxa thrive more in an anaerobic environment with depleted resources. Such taxa included *Thermincola* (c. Clostridia) and *Candidatus Solibacter*, which were only identified in high proportion in samples incubated for 10 or more days (Figure 4).

### 3.3 The abundance of nitrogen, sulfur, and methane metabolism genes

PCR assay results are shown in Table 1, where genes related to denitrification, DNRA, sulfate reduction, and methanogenesis were amplified. In contrast, we did not observe copies or amplification of anammox genes. The qPCR results corroborated previous study by Adyasari et al. (2020) which predicted the occurrence of denitrification and DNRA. However, qPCR for denitrifying genes showed higher abundance of such genes (ranging between 3.10^6^ – 3.10^7^ gene copies/gr sediment [Figure 5]) compared to less than 10^2^ gene copies/gr sediment for DNRA. The lowest and highest average abundances of denitrification genes were observed in control and groundwater-spiked samples, respectively. Natural seawater-spiked incubations exhibited the lowest denitrification abundance between samples with different type of saline solution.

**Figure 5.**
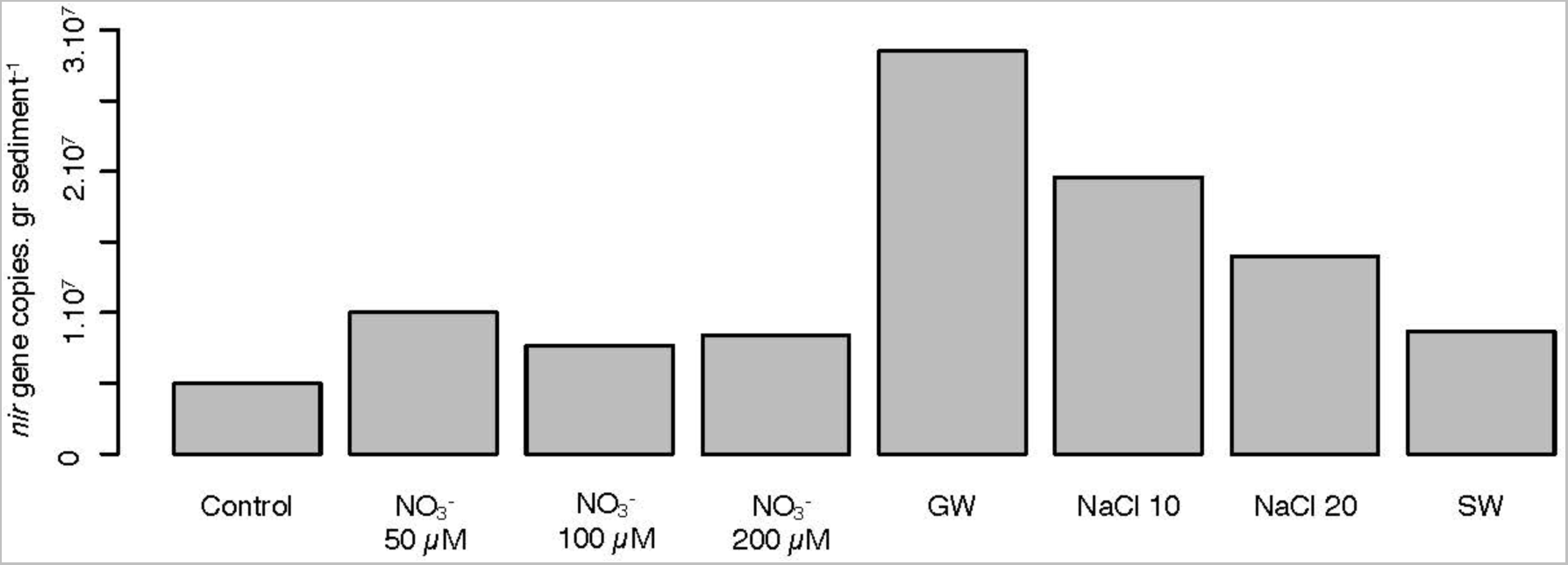
Quantitative PCR results for denitrifying genes in different treatment group. GW= groundwater, SW = seawater.

**Table 1.**
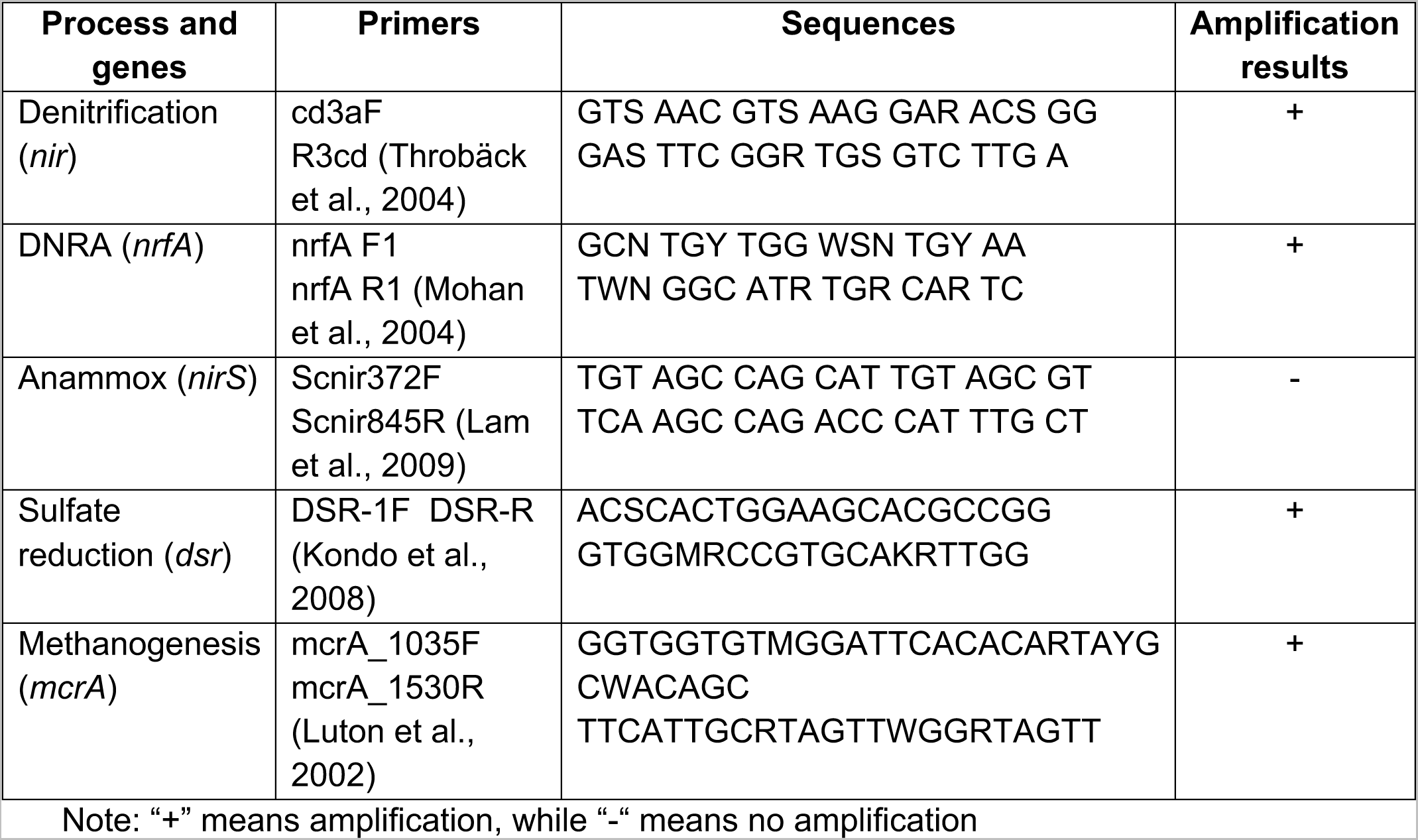
List of biogeochemical processes, primers, sequences, and amplification results of PCR assay

### 3.4 Linkage between microbial communities and DOM molecular compounds

To examine microbial roles in DOM processing, we performed Spearman correlations between taxonomic data on the class level with DOM formulae generated from FTICRMS (Table S1). Out of 79 classes observed across all samples, eight displayed significant correlations with certain DOM compounds (Figure 6). Van Krevelen diagrams showed groups such as Bacilli, Methylomirabilia, SRB (Desulfarculia, Desulfomonilia, and Desulfitobacteriia), and fermenters (Spirochaeta) were negatively correlated with aromatic compounds. ASVs belonging to Campylobacteria were associated with both labile (unsaturated) and less labile (highly unsaturated) compounds.

**Figure 6.**
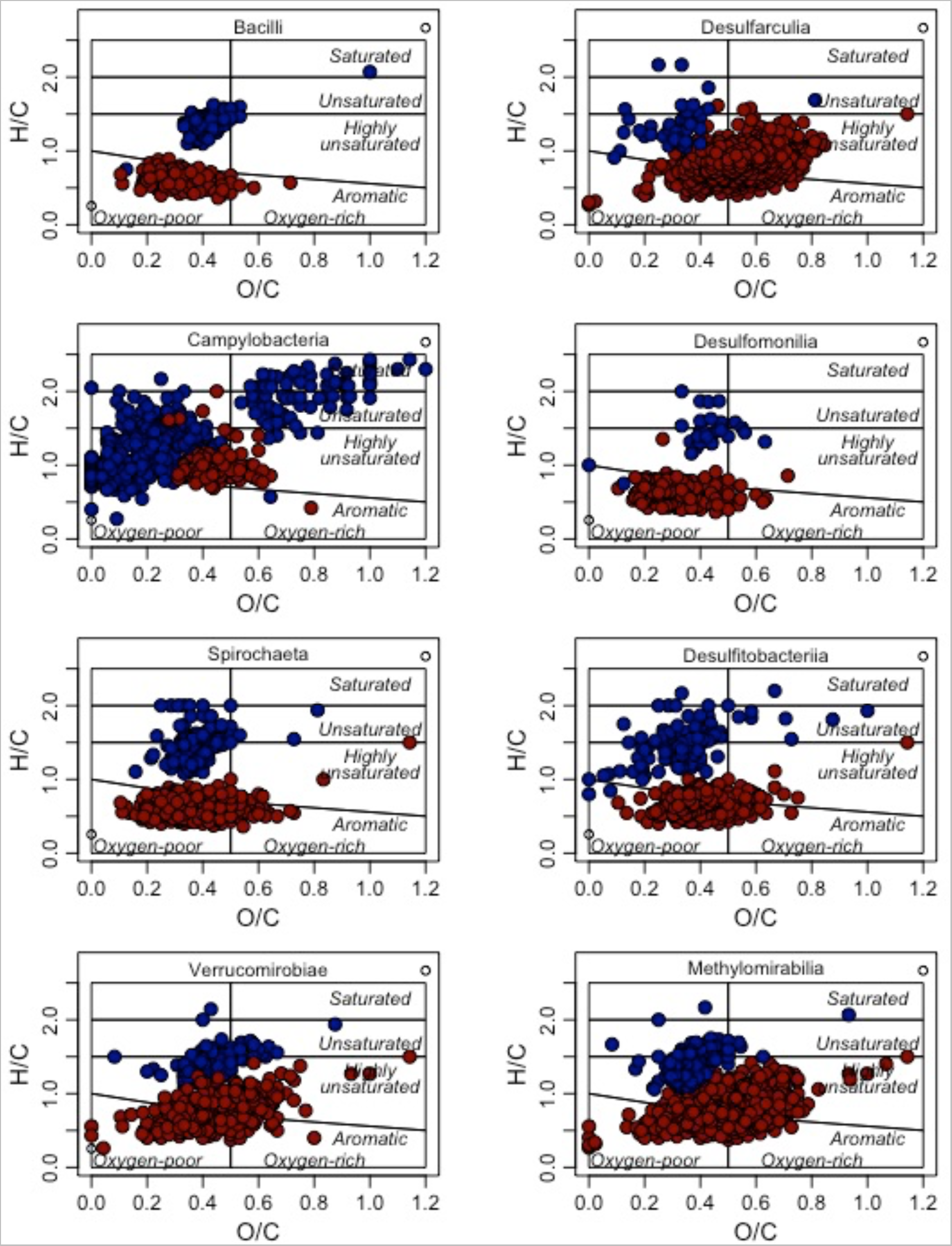
Van Krevelen plots of compounds significantly correlating with microbial classes (Spearman rank test, p < 0.05). Blue and red color indicates positive (ρ > 0.5) and negative (ρ < −0.5) correlations, respectively.

### 3.5 Predicted microbial metabolic pathways

In this study, PICRUST2 was employed to predict metabolic functions based on 16S rRNA gene sequencing data. The following pathways were described according to their name in the MetaCyc database (Caspi et al., 2016). Fermentation, sulfur, and methane metabolism were observed in all samples (Figure 7). Methanogenesis from acetate (METH-ACETATE-PWY) was more dominantly predicted than a similar process using H_2_ and CO_2_ (METHANOGENESIS-PWY). This finding complemented the fermentation results, where some highly predicted processes had acetate as the final products (e.g., PWY-6588, PWY-6590, PWY-5100). Methylotrophic pathway (PWY-7616), i.e., methane or methanol utilization to CO_2_, was also predicted but in lower abundance than methanogenesis.

**Figure 7.**
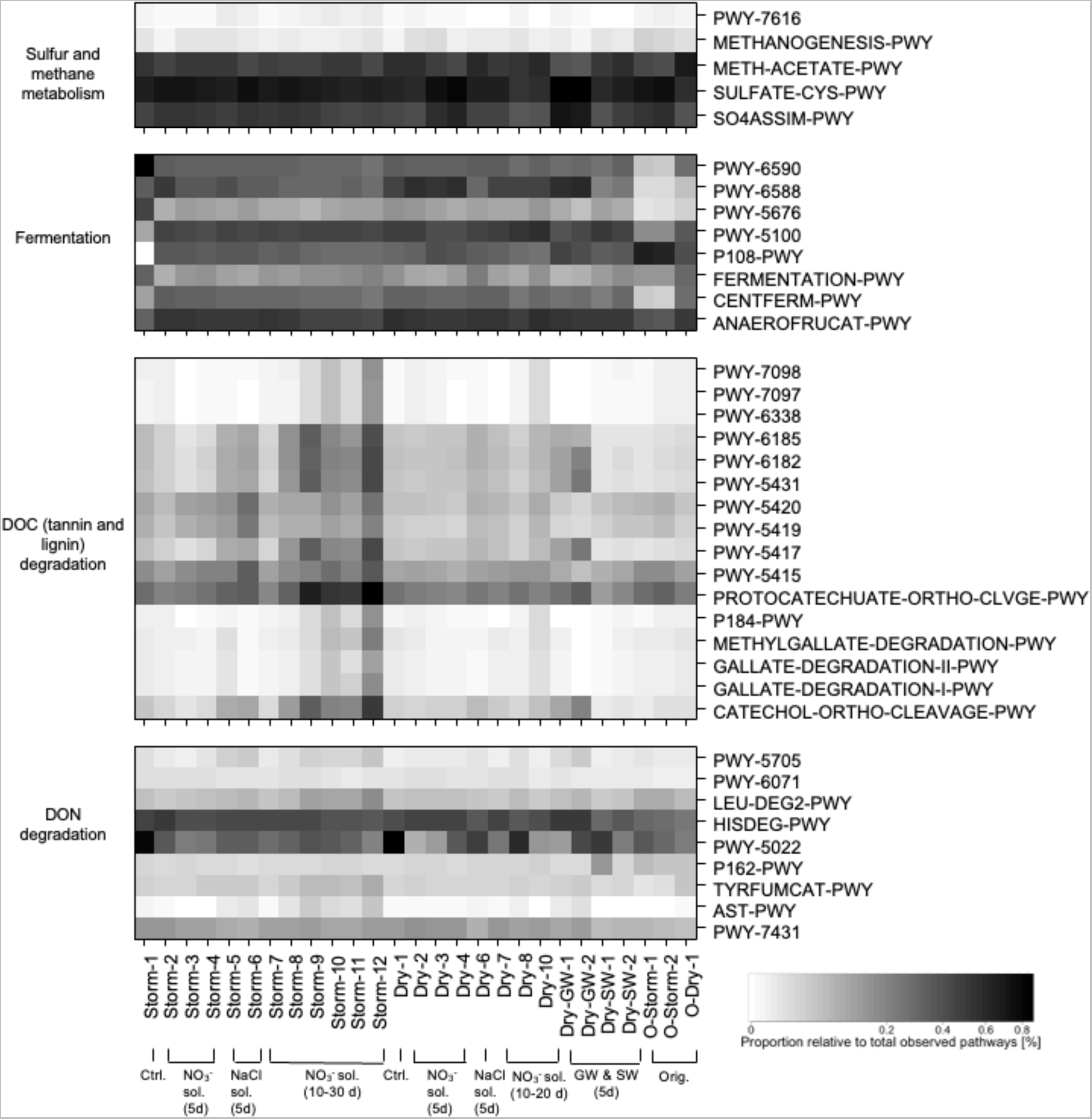
Heatmap of the proportion of predicted pathways related to sulfur and methane metabolism, fermentation, DOC, and DON degradation based on PICRUST2. The description of each pathway can be seen in Table S5. Abbreviations “Ctrl.”, “GW”, “SW”, and “Orig.” are control, groundwater, seawater, and original sediment incubations, respectively.

Related to plant material degradation, PICRUST2 predicted microbial communities’ role in degrading aromatic (PWY-5431), catechol and methyl catechol (PWY-6185, PWY-5417, PWY-5419, PWY-5420, CATECHOL-ORTHO-CLEAVAGE-PWY), gallate (GALLATE-DEGRADATION-I-PWY), methyl gallate (METHYLGALLATE-DEGRADATION-PWY), vanillin and vanillate (PWY-7098, PWY-7097, PWY-6338), and protocatechuate (PROTOCATECHUATE-ORTHO-CLEAVAGE-PWY) across all samples. The highest abundance of these degradation pathways was observed in the longest incubation samples in each batch (i.e., Storm-12 and Dry-9). In terms of DON degradation, PICRUST2 result indicated that L-histidine degradation (HISDEG-PWY), 4-aminobutyrate degradation (PWY-5022), and aromatic biogenic amine degradation (PWY-7431) pathways were the most dominant processes across all samples. However, we did not observe specific patterns of these pathways associated with experimental treatments.

## 4. Discussion

### 4.1 Core communities and rare biosphere associated with plant-derived organic matter degradation

In this study, we conducted microcosm experiments to elucidate the microbial responses to environmental changes in organic-rich subterranean estuaries. We observed that the original sediments (prior to incubation) were dominated by Clostridia, Bacilli, and Desulfobacteria, with other classes such as Gammaproteobacteria, Alphaproteobacteria, Anaerolinae, Desulfitobacteriia, and Desulfobaccia comprising up to 70% of the identified microbial community composition. Clostridia and Anaerolinae are both obligate anaerobic taxa and usually found in deep, sulfidic layers of STEs (Ruiz-González et al., 2022), while Gammaproteobacteria, Alphaproteobacteria, and SRB (e.g., Desulfobacteria, Desulfitobacteriia, and Desulfobaccia) have been reported in other studies investigating sedimentary microbial STEs (Degenhardt et al., 2020; Hong et al., 2019). The SRB, Anaerolinae, and Thermodesulfovibrionia were also observed in a previous field study at the same site, where porewater was collected and analyzed instead of sediments (Adyasari et al., 2020). A certain overlap between free-living and particle-attached bacteria in environments is common (Urvoy et al., 2022). In our case, the notable difference between free-living and particle-attached bacteria is the relative proportion of Clostridia, which only constituted 1-5% in porewater samples; however, their relative proportion increased to 35% in the sediment samples. Their characteristic as soil bacteria explained their prevalence in STE sediment over porewater (Wells and Wilkins, 1996).

We observed an altered microbial community composition between original and treated sediments. Our results indicated that the core communities survived environmental changes while in contrast, the less adaptable taxa were extinguished. A previous study reported that Leptospirae, one of the taxa not found after the incubation, had difficulties multiplying following a microcosm experiment (Casanovas-Massana et al., 2018). Another taxon, Omnitrophia, is known for its host-dependent lifestyle (Seymour et al., 2022). It is likely that the altered environment during our incubation removed such hosts and lessened their survivability.

After incubations, Clostridia and Bacilli emerged as the most prevalent taxa (Figure 4). This finding is unsurprising, considering both classes are known to withstand abrupt change or harsh environments; they are motile, adaptable to pH and oxygen availability, tolerate moderate salinity, and can form endospores as a response to nutritional stress (Eichenberger, 2013; Errington, 2003). Gammaproteobacteria, the most phylogenetically rich taxon of the prokaryotes, comprised several highly adaptable, cosmopolitan genera (e.g., *Acinetobacter* and *Pseudomonas*) (Brenner et al., 2005). SRB and methanotrophic bacteria from Alphaproteobacteria and Gammaproteobacteria (e.g., *Methylobacter* and *Methyloceanobacter*) also flourished in treated sediments because of reducing conditions during incubation. The relative proportion of iron reducers, represented by ASVs from the family Geobacteraceae (Röling, 2014), was stable across all samples (i.e., between 1-2%). While their abundance was not as dominant as SRB, their identification was likely considering STE sediments contain notable Fe concentrations (Lamore, 2020). Overall, core communities in sediments consisted of a mixture of terrestrial and marine sedimentary bacteria, consistent with STE’s characteristics as a land-ocean interface. The resiliency of the core sedimentary community was also found in other STEs (Calvo-Martin et al., 2022; Degenhardt et al., 2020). These seasonal studies concluded that, notwithstanding slight fluctuations in relative proportion, the dominant sedimentary communities remain largely stable despite salinity and temperature variability throughout the year.

PICRUST2 predicted the abundance of pathways related to vascular plant organic matter degradation across all samples. Some bacteria can initiate tannin and lignin degradation by hydrolyzing these complex organic compounds using extracellular enzymes and yielding sufficiently small metabolites that can be processed further. Some of the metabolites and final products of plant organic matter predicted included (1) gallate, methyl gallate, catechol, methyl catechol, vanillin, vanillate, and protocatechuate, i.e., metabolites and final products from lignin and tannin degradation (Bhat et al., 1998; Bugg et al., 2011; Sharma et al., 1998), (2) salicylate, a plant hormone (Raskin, 1992), and (3) toluene, i.e., components produced and emitted by plants (Parales et al., 2008). The strongest prediction across all samples was exhibited by protocatechuate degradation, indicating that a major portion of lignin and tannin have been biologically degraded, as this pathway is the last step in the microbial lignin utilization process (Bugg et al., 2011). We also crosschecked the occurrence of these chemical compounds to FTICRMS results (Table S2) and found their formulae across all samples. However, it should be noted that formula abundance does not equal to target compounds; therefore, these association should be interpreted with cautions. Furthermore, studies linked the tannin and lignin degradation pathways to Gammaproteobacteria (e.g., *Acinetobacter*), Alphaproteobacteria, and Bacilli (Bugg et al., 2011; Olsson, 2016), some of the core taxa in the treated sediments. Clostridia, Bacilli, and other bacteria with high relative proportion found in this study (e.g., Verrucomicrobiae) possess polymer hydrolases responsible for breaking down particulate or high-molecular weight organic carbon into smaller compounds (Abd-Elsalam and El-Hanafy, 2009; Gihring et al., 2009; Matsushita and Okabe, 2001). For Bacilli, Spearman correlation analysis result supported this finding: a negative and positive correlation was observed between this taxon, aromatic, and highly unsaturated compounds, respectively (Figure 6). This result implies their role in catabolizing high molecular weight, aromatic compounds and producing more resistant, lower molecular weight compound in DOM degradation process.

The metabolites from hydrolysis is usually consumed by the fermenters. Clostridia and Anaerolinae, for instance, belong to this functional group. The products of fermentation, e.g., fatty or carboxylic acids, can be further utilized by denitrifiers, SRB, and methanogens, which were abundant in treated sediments. The presence of active fermentation, sulfate reduction, and methanogenesis was corroborated by PCR (Table 1) and PICRUST2 results (Figure 7), which showed amplifications and predictions of these processes. The correlation between SRB and fermenters in Figure 6 also suggests their role in degrading higher molecular weight, aromatic compounds to lower molecular weights, highly unsaturated or unsaturated compounds.

The complete syntrophic association between POC degraders, fermenters, and methanogens to transform terrigenous, plant-derived DOM compounds into intermediate and final products has been observed elsewhere (Field and Lettinga, 1992). In the last step, methanogens can remove organic acids and hydrogens, which create thermodynamically favorable conditions for the cleavage of the aromatic ring (Bhat et al., 1998). Interestingly, the relative proportion of methanogenic bacteria and archaea was considerably low across all samples (less than 1% of entire communities). The disproportionate functional role of rare taxa, i.e., they have been shown to support specific ecosystem processes more strongly than expected from their abundance, is a widespread phenomenon and has also been observed in other STEs (Héry et al., 2014; Ruiz-González et al., 2022).

We also used PICRUST2 to predict DON degradation pathways consisting of amino acids, amines, and polyamines across all samples (Figure 7). These DON catabolic pathways are possessed by a wide range of bacteria (including Clostridia and Bacilli) and generated NH_4_^+^ as a metabolite or final product (Borek and Waelsch, 1953). The most robust prediction among DON catabolic pathways was exhibited by the 4-aminobutyrate degradation pathway (PWY-5022), a compound present in the environment as a product of plant and animal tissue decay (Bouché and Fromm, 2004). This pathway is widely possessed by bacteria in phylum Firmicutes (including Clostridia and Bacilli) to produce NH_4_^+^ and acetate. Again, this finding corroborates with PICRUST2 results on methanogenesis prediction, where the acetogenic pathway is preferred and likely to occur in sediments. The abundance of DON catabolic pathways and the absence of several NH_4_^+^ utilization pathways (e.g., anammox [Table 1], nitrification [due to the hypoxic conditions]) across all samples indicated active internal cycling of N that accumulates NH_4_^+^ in STE sediments. When groundwater or seawater transits through STE, the ionic exchange between sediment surface and water creates a condition where NH_4_^+^ desorbs to water, consequently creating NH_4_^+^-rich SGD in Mobile Bay, as seen in Montiel et al. (2019b).

In general, our results indicated that microbial metabolic pathways, specifically DOC and DON degradation, were not overly sensitive to environmental changes when (1) most of the processes were performed by resilient, core STE communities and thus promoted functional redundancy, and (2) there was an abundance of electron donors (i.e., high DOC concentration) and recipients (i.e., NO_3_^−^ or SO_4_^2−^) for such metabolisms to occur.

### 4.2 Seasonal sampling, groundwater, and seawater addition influence the dynamics of less prevalent taxa

In this study, we observed that seasonal sediment collection resulted in slight difference of particle size distribution; however, the microbial community composition between sediment batch was almost similar. At the genus level, the most notable difference between the two seasonal samples was the identification of *Pseudarcobacter* (c. Campylobacteria) in high abundance in the Storm batch. However, their relative proportion was significantly lower in Dry samples. *Pseudarcobacter* was likely dispersed to STEs as the soil particles were resuspended during heavy rain, happened more often during storm season. They may have originated from marine biota and sediments, from which species of *Pseudarcobacter* have been isolated (Pérez-Cataluña et al., 2018). This taxon exhibited the most positive association with the labile DOM pool identified by FTICRMS (Figure 6), suggesting their role in processing these organic compounds in Storm samples. The capability of Campylobacteria to degrade organic constituents was also observed in another study by Feng et al. (2022), where ASVs associated with this class were found to be responsible for degrading secondary metabolites of kelp detritus in coastal environments.

Contrary to *Pseudarcobacter, Exiguobacterium* (c. Clostridia) was more ubiquitous in Dry than Storm samples. This genus is known as NO_3_^−^ reducer and produces enzymes such as alkaline proteases, phosphatase, dehydrogenase, and decarboxylase, which are involved in the biodegradation of several complex compounds (Pandey, 2020). This result agreed with the sedimentary properties of Dry samples, which exhibited slightly higher aromatic and highly unsaturated compounds than Storm samples.

In experiments with groundwater and seawater input, we observed taxa in treated sediments that likely originated from those endmembers in Mobile Bay. For instance, groundwater-spiked samples displayed a higher relative proportion of *Fictibacillus* (c. Bacilli) than other samples (Figure 4). The anaerobic strains of this taxon have been isolated from freshwater (Glaeser et al., 2013; Pal et al., 2018). They plausibly originated from groundwater and, considering their spore-forming capability, attached to sediment particles during incubations where they were identified during analysis. A similar attachment pattern was also displayed by *CL500-29 marine group, Ilumatobacter*, and *Cyanobium PCC-6307*—all were taxa ubiquitous in Mobile Bay seawater (Adyasari et al., 2020).

In general, the introduction of allochthonous microorganisms to subsurface environments during salinization, heavy precipitation, recharge-discharge episodes, or storm events has been observed elsewhere (Retter et al., 2021). This phenomenon may invade or reassemble indigenous microbial communities, subsequently altering local community composition and ecosystem functioning. Extreme cases occurred in geologically fractured areas, where an altered anchialine microbial community was observed when decreasing groundwater table pulled seawater inland (Héry et al., 2014). Our study indicates that such mobilization possibly occurs in coastal sedimentary aquifers but to a lesser extent, as a result of (1) unconsolidated sediments restraining the microbial movement (i.e., environmental filtering), and (2) the resilience of core communities preventing significant alteration of overall microbial community composition.

### 4.3 Denitrification was susceptible to groundwater and seawater addition

The findings from geochemical measurements and qPCR assays indicated an active denitrification process during incubation. However, this study did not observe significant changes in microbial community composition across samples incubated with artificial NO_3_^−^ and groundwater solution. The consistent taxonomic assemblages under different NO_3_^−^ concentrations could stem from (1) nitrite reductase genes being widespread among bacteria, including those identified in this study (Tiedje, 1988), and (2) the abundance of electron donors (i.e., DOC concentration up to 3,000 µM [Table S1]) creating a condition where NO_3_^−^ was not a limiting electron acceptor in Mobile Bay STE. However, as NO_3_^−^ generates the highest energy out of other electron acceptors in anaerobic redox reactions, microorganisms prefer to utilize it in their metabolic activities. This notion is supported by the outcomes of the qPCR assay (Figure 5). The result demonstrated a higher abundance of denitrifying genes was found in samples treated with NO_3_^−^ solution, NaCl solution, groundwater, and seawater compared to the control, which was only infused with Nanopure water (NO_3_^−^ = 0 µM). Between samples with different NO_3_^−^ inputs, NO_3_^−^ concentrations in Dry-4 (NO_3_^−^ = 200 µM) and Dry-GW-1-2 (NO_3_^−^ = 210 µM) were similar; however, the quantified genes were higher in the latter. This finding implies the possibility of (i) existing components in groundwater, that were not available in artificial NO_3_^−^ solutions and activated denitrification, or (ii) the qPCR assay quantified inactive denitrification genes from groundwater collected from the field. Furthermore, this study’s denitrifying gene quantification result is comparable to or higher than other STE studies, where denitrification is considered a major pathway of NO_3_^−^ removal (Jiao et al., 2018; Wu et al., 2021).

In incubations with different saline solutions, we found that denitrifying genes were more abundant in sediments incubated with artificial NaCl (NO_3_^−^ = 50 µM) than in sediments mixed with seawater solution (NO_3_^−^ = 2 µM). Dry/Storm-6 samples (salinity = 20) also exhibited a higher abundance of denitrifying genes even though they had higher salinity than seawater (salinity = 15). This result indicates that chemical components naturally occurring in seawater may have been the factors negatively affecting the denitrification capability of microorganisms. Indeed, cations other than Na (e.g., Mg^2+^, Ca^2+^) and sulfide have been reported as adversely impacting denitrifying microorganisms and lowering denitrification potentials in coastal sediments (Aelion and Warttinger, 2010; Macêdo et al., 2019; Murphy et al., 2020).

Overall, our study suggests that denitrification is sensitive to environmental changes such as salinization and NO_3_^−^ inputs. STE salinization of local aquifers results from (1) sea level rise, and (2) excessive groundwater withdrawal, while the dynamics of groundwater NO_3_^−^ concentrations are affected by the land use management and the level of anthropogenic activities. While NO_3_^−^ was successfully removed from all porewater in this particular study, another study by Lamore (2020), which investigated denitrification capability in sediments collected at the same site, indicated that the sediments’ capacity to attenuate NO_3_^−^ decreases as the load increases. The collective findings imply that groundwater management, i.e., reducing groundwater NO_3_^−^ concentrations and groundwater extraction rates, should be priorities in conserving the denitrification capacities of STEs.

### 4.4 Long-term incubations favor taxa with specific survival capability

This study observed a specific pattern where the more robust predictions of DOC degradation occurred in porewater incubated > 5 days, with the highest prediction from 35 days of incubation. This finding agrees with the degradation state in such samples (Figure 3), indicating that a long incubation period is analogous to the degradation rate. This outcome implies that, given sufficient incubation time (i.e., residence time in STE), microbial communities can utilize less labile DOM compounds and therefore promote the re-release of buried solid phase DOM into the active C cycle.

We observed an increase in nASV and diversity in the long-term incubations; however, rare taxa decreased in abundance with increased incubation time (Figure 2). This finding could stem from different factors, e.g., surviving taxonomic groups were more capable of accumulating and storing specific nutrients, blocking access to the competitor’s favorable habitats, having more motility than the competitors, or forming spores or other protective extracellular enzymes (Hibbing et al., 2010). That dormancy contributes to microbial diversity has been observed elsewhere, where dormant bacteria accounted for 40% of taxon richness in nutrient-poor systems (Jones and Lennon, 2010). Two of the characteristics mentioned above were exhibited by *Thermincola* (c. Clostridia), which was not identified in the original sediments but who accounted for up to 5% of the microbial community in long-term incubations. *Thermincola* is a motile, spore-forming bacterium that can grow chemolithoautotrophically (Sokolova et al., 2005; Zavarzina et al., 2007), which explains its dormancy in non-incubated sediments and short-term incubations, but high presence in long-term incubations. *Candidatus Solibacter* (c. Acidobacteriae), another genus that thrived in long-term incubations, is equipped with a large number of anion:cation symporters and engages in biofilm production (Ward et al., 2009). The same study found that these functions benefit *C. Solibacter*, as they increase their ability to survive in low-nutrient environments and under environmental stress conditions. In addition to these two genera, ASVs related to SRB and methanotrophic bacteria also had an increased abundance in incubations > 5 days, attributed to their capability to utilize fermentation products or a variety of multi-carbon compounds (Takeuchi et al., 2014).

Linking these microcosm data to actual site conditions, our results indicated that prolonged groundwater transit time in STE during an extended drought period alters microbial diversity on a local scale, causing mortality to the less adaptable taxonomic groups and preferring the survival of opportunistic bacteria and taxonomic groups associated with sulfate reducer or methane oxidation. The sensitivity of rare biosphere towards changing incubation times is a particular concern, considering their immense role in the syntrophic consortium of organic matter degradation, as discussed in section 4.1. The loss of partners in such consortia may have severe consequences for ecosystem functioning, unless the species loss is compensated by other taxonomic groups that also thrive in the changing environments (i.e., functional redundancy capability).

## Conclusion

In this controlled laboratory study, geochemical and metagenomic approaches were employed to elucidate the microbial response to sudden environmental changes and its implications for metabolic pathways, including denitrification and organic matter degradation. We found that most STE sediments consist of resilient, core taxonomic groups that contribute to C and N cycles in STE. Seasonal sampling, incubation times, and groundwater and seawater addition affected the relative proportion of less prevalent taxonomic groups. The organic-rich, anaerobic conditions of STE sediments supported a syntrophic consortium of POC degraders, fermenters, and acetogenic methanogens catabolizing plant-derived organic compounds. In terms of metabolic pathways, DOC and DON degradation predictions were not significantly affected by such changes, while denitrification was found to be susceptible to seawater intrusion. The results of this study highlight the sensitivity of rare biosphere (i.e., methanogenic archaea), denitrification processes, and less adaptable microbial assemblages to changes in climate, land, or water use. A particular focus should be applied to reducing groundwater withdrawal rates and NO_3_^−^ concentrations in the terrestrial aquifer to maximize the denitrification capacity of STE sediments. In addition, the results of this study can be used as an input for ecological modeling or as a base for future studies to investigate whether the results translate quantitatively to ecosystem-scale biogeochemistry.

## Acknowledgments

The authors wish to thank the student assistants during field and laboratory work, Stephen Anderson, Justin McCleskey, Julian Wielkens, and Charles Drumm. D.A. is funded by the German Research Foundation (DFG) through the Walter Benjamin Fellowship (No. 446330207) and the NSF EPSCoR RII Track-2 FEC: Emergent Polymer Sensing Technologies for Gulf Coast Water Quality Monitoring granted to NTD. H.W. is grateful to Ina Ulber, Matthias Friebe, and Katrin Klaproth for assistance in DOC and DOM analyses, and acknowledges funding from the DFG research unit “DynaDeep” (FOR 5094, WA3067/3-1).

